# Predicted meta-omics: a potential solution to multi-omics data scarcity in microbiome studies

**DOI:** 10.1101/2024.11.04.621857

**Authors:** Bianca-Maria Cosma, Stephanie Pillay, David Calderón-Franco, Thomas Abeel

## Abstract

Imbalances in the gut microbiome have been linked to conditions such as inflammatory bowel disease, diabetes, and cancer. While metagenomics and amplicon sequencing are commonly used to study the microbiome, they do not capture all layers of microbial functions. Other meta-omics data can provide more insights, but these are more costly and laborious to procure. The growing availability of paired meta-omics data offers an opportunity to develop machine learning models that can infer connections between metagenomics data and other forms of meta-omics data, enabling the prediction of these other forms of meta-omics data from metagenomics. We evaluated several machine learning models for predicting meta-omics features from various meta-omics inputs. Simpler architectures such as elastic net regression and random forests generated reliable predictions of transcript and metabolite abundances, with correlations of up to 0.77 and 0.74, respectively, but predicting protein profiles was more challenging. We also identified a core set of well-predicted features for each meta-omics output type, and showed that multi-output regression neural networks performed similarly when trained using fewer output features. Lastly, our experiments demonstrated that predicted features can be used for the downstream task of inflammatory bowel disease classification, with performance comparable to that of experimental data.

## 1. Introduction

The human microbiome directly and indirectly engages with various physiological subsystems, including the nervous, gastrointestinal, cardiovascular, and immune systems. Research has shown that imbalances in the human microbiome, commonly referred to as dysbiosis, are associated with the onset and progression of various health conditions. For instance, the composition of the gut microbiome, along with its associated metabolites, was found to be significantly different between healthy individuals and those suffering from IBD, type I and II diabetes, cardiovascular disease, as well as mental health disorders such as depression and anxiety.^1–3^ Dysbiosis in the vaginal microbiome has been linked to cervical cancer, as it can affect the development and advancement of HPV (human papillomavirus) infection.^4^ Beyond the human microbiome, recent literature also highlights the importance of investigating microbial communities in non-clinical sectors, with applications ranging from surveillance of antibiotic resistance genes to the study of greenhouse gases.^5,6^

Microbial communities can be characterized across various layers of functional activity, with state-of-the-art meta-omics technologies such as metagenomics (mGx), metatranscriptomics (mTx), metaproteomics (mPx), and metabolomics (mBx), among others. To obtain a complete picture of the microbiome, each sample should be characterized using all of these meta-omics modalities. Although the metagenome encodes the functional potential of a microbial community, the presence of genes is not synonymous with active transcription into mRNA, and the latter is not always translated into active proteins. Additionally, even though some associations between microbes and metabolites are known, these can be ambiguous and inconclusive, due to the fact that two microbes may produce the same metabolite, or that some metabolites may only be produced under certain conditions.^7^ The importance of data accessibility across diverse meta-omics layers is not only emphasized in experimental research, but also in the development of machine learning models capable of performing a wide range of predictive tasks, including disease detection.^8–14^

However, measuring meta-omics data comes with many challenges, including significant costs and reliability issues.^15^ DNA sequencing data, whether in the form of amplicon or shotgun metagenome sequencing, is currently the most accessible option, due to its lower cost and the higher reliability provided by next-generation sequencing. At the same time, paired multi-meta-omics data is becoming increasingly available, through initiatives such as the Integrative Human Microbiome Project.^16^ This availability of microbiome data across multiple meta-omics layers presents an opportunity to develop machine learning models capable of inferring connections between metagenomics data and other forms of meta-omics data, with the eventual goal of predicting the latter from the former.

Several studies have already described the use of machine learning to predict metabolite abundances in microbiome samples, starting with features derived from metagenomics data. Some model architectures that have been proposed include MelonnPan (elastic net regression), SparseNED (sparse neural encoder-decoder), MiMeNet (multilayer perceptron), mNODE (neural ordinary differential equations), MMINP (two-way orthogonal partial least squares (O2-PLS)) and LOCATE (neural network).^17–22^ However, the prediction of other meta-omics modalities, in addition to metabolomics, has not been investigated.

In this manuscript, we propose a novel application of metabolite prediction models to infer the abundance of microbial transcripts and proteins. To that end, we perform a benchmark of multiple machine learning models on the task of metatranscriptomics, metaproteomics and metabolomics prediction, from various meta-omics inputs. We show that these models can generalize to multiple input-output combinations of meta-omics modalities, generating reliable predictions for a core set of transcripts, proteins and metabolites. To demonstrate the utility of such prediction models in microbiome research, we highlight an application of predicted meta-omics data for IBD diagnosis. Our methodology provides a starting point for further development of machine learning pipelines that can perform integration and prediction across multiple meta-omics layers, with applications in the diagnosis and treatment of microbiome-associated conditions.

## 2. Materials and methods

### 2.1. Analysis of feature filtering for meta-omics prediction

To determine the degree to which sparse meta-omics features should be filtered out, we performed initial benchmarking of our experimental pipeline on three datasets containing paired metagenomics and metabolomics data (Supplementary Table S1): Franzosa et al.^13^ (inflammatory bowel disease), Wang et al.^23^ (end-stage renal disease) and Yachida et al.^24^ (colorectal cancer). The latter two datasets were downloaded from The Curated Gut Microbiome Metabolome Data Resource, release v2.1.0.^25^ The IBD dataset was downloaded from the paper’s supplement. The IBD dataset contains metagenomics data in the form of gene families, while the other two contain taxonomic profiles at the species level. We note that MelonnPan was originally trained and tested on the same IBD dataset.^17^

We evaluated two approaches for filtering out low-abundance features (species, genes, and metabolites):

- **strict filtering:** similarly to Mallick et al.^17^, we retained only those features with at least 0.01% abundance in more than 10% of samples. We additionally filtered out features with less than 0.0001% abundance in more than 10% of samples. Features with more than 95% zeros were also filtered out across all feature types.
- **lenient filtering:** we retained only features with at least 0.005% abundance in more than 10% of samples. Features with more than 95% zeros were filtered out.

For each filtering method, we ran MelonnPan on all three datasets, with default settings, to predict metabolite abundances from metagenomics data. All benchmarking was performed on separate test sets. The data collected by Franzosa et al.^13^ included an independently sampled validation cohort, which we used as a test set. We split the two remaining datasets into a training set and a test set, with a ratio of 80% to 20%.

The predicted data, along with the input metagenomics and experimental metabolomics data, were subsequently used to classify disease. To that end, we trained 10 random forest classifiers, initialized with different random seeds (the same ones shown in Supplementary Table S3) to predict phenotypes specific to each dataset. We used scikit-learn’s RandomForestClassifier (v1.4.1.post1), with default parameters.

### 2.2. Data processing on IBDMDB

#### Gut microbiome meta-omics data

We downloaded pre-processed metagenomics, metatranscriptomics, metaproteomics and metabolomics data from IBDMDB (Inflammatory Bowel Disease Multi’omics Database), which was assembled as part of the Integrative Human Microbiome Project.^26^ The dataset contains longitudinal samples from 132 subjects, including a control group, as well as patients diagnosed with ulcerative colitis (UC) or Crohn’s disease (CD). Download links and dates are recorded in Supplementary Table S2. Before feature filtering, all meta-omics abundance profiles were normalized, such that feature values per sample sum up to 1. We used gene, transcript and protein abundance profiles annotated using Enzyme Commission numbers (ECs). As this data was originally stratified, we summed up ECs across taxonomic groupings to reduce dimensionality and sparsity. Additional experiments supporting all of our main results were performed using pathways and species abundances derived from mGx data, as shown in some of the supplementary results (Supplementary Tables S3 and S7). For mBx data, we retained one LC-MS technology, namely C18 negative (C18-neg).

#### Imputation of zeros

To enable log transformation of features at a later stage in our pipeline, we also generated versions of these datasets with imputed zeros. For mGx and mTx data, we added *ϵ* = 1e-6 to all abundances, which is less than all other non-zero values in the matrices, while for mPx and mBx, which were available as count data, we added a pseudocount.

#### Paired meta-omics datasets

We generated paired meta-omics datasets for multiple input-output combinations of meta-omics modalities. Experiments were set up as follows:

- predicting transcripts (mTx) from genes (mGx);
- predicting proteins (mPx) from genes (mGx) and transcripts (mTx);
- and, lastly, predicting metabolites (mBx) from genes (mGx), transcripts (mTx) and proteins (mPx).

In addition, we also predicted mPx and mBx data from multi-omics input, obtained as combinations of single-omics input, constructed using standard feature concatenation. In total, we analyzed results from 11 different input-output combinations of meta-omics modalities. Supplementary experiments were performed for a total of 32 models, including input data types represented as taxonomic profiles and pathways extracted from metagenomics data. A full overview of all paired datasets is provided in Supplementary Table S3.

#### Feature filtering

Across all datasets, we applied the lenient feature filtering approach previously described in Section 2.1.

#### Data transformations

Following feature filtering, each sample was normalized and the data was transformed to account for compositionality, sparsity, and feature scaling. To that end, we compared two standard transformations for compositional data, namely the centered log ratio (CLR) transformation, which requires the imputation of zeros, and the arcsin square root transformation, which also works on non-imputed data. The CLR transformation of a sample *x* ϵ ℝ^*D*^, with sum of elements 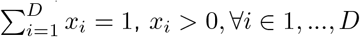, and *g*(*x*) defined as the geometric mean of *x*, is given by:

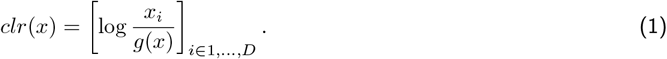

For *x* ϵ ℝ^*D*^, with sum of elements 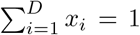 and 0 ≤ *x*_*i*_ ≤ 1, ∀*i* ϵ 1, …, *D*, the arcsin transformation is as follows:

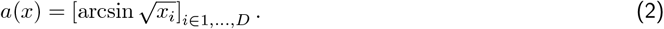

As initial experiments showed that MelonnPan performed best among all benchmarked models, we also tested the quantile transformation implemented for this model, which maps normalized features to the quantiles of a normal distribution. This transformation was shown to improve the predictive power of standard regression models and neural networks.^27,28^ Although Mallick et al.^17^ only apply this transformation to the input features, we transformed the output features as well, to preserve consistency with other transformations that we benchmarked. This was implemented using scikit-learn’s QuantileTransformer (v1.4.1post1), with the output distribution set to “normal”.

### 2.3. Training and evaluation of meta-omics prediction models

#### Training and testing partitions

To make up for the lack of an independently sampled test set and provide a fair evaluation, we generated 10 train/test splits of each paired dataset, based on a fixed set of random seeds, with a ratio of 80% to 20% (Supplementary Table S3). To reduce overfitting, we performed each split on patients instead of samples, such that samples belonging to the same patient would not be present in both the training and test sets. Each partition was stratified, preserving the proportion of classes (UC, CD, and healthy control (HC)) between the training and test samples. Aside from the paired datasets, we applied the same procedure for three non-paired datasets containing mTx, mPx and mBx data (see Supplementary Table S4), which are later used in benchmarking IBD classifiers (Section 2.4).

#### Benchmarking of cross-omics regression models

We benchmarked four models and a baseline on several cross-omics prediction tasks. From the literature, we selected MelonnPan (elastic net regression), SparseNED (sparse neural encoder-decoder) and MiMeNet (feed-forward neural network).^17–19^ These architectures were all originally designed to predict metabolite abundances from metagenomics. All models were run with default parameters, except for MiMeNet, where some parameters were changed to reduce runtime (see Supplementary Note 1). Each model was trained and tested using different data transformations (see Supplementary Table S5). We also trained a deep neural network (Deep NN), with data augmentation (Supplementary Note A.1.1), and a RandomForestRegressor baseline (scikit-learn v1.4.1.post1), initialized with default parameters and a random seed equal to 42. For more details regarding the network architecture, as well as the loss used for training, see Section A.1 of the Supplementary Methods, particularly Subsections A.1.2 and A.1.3. Hyperparameter tuning for the neural network is also recorded in Supplementary Table S9.

#### Model evaluation

We evaluated all models on each independent test set by comparing predicted features (transcripts, proteins, and metabolites) with the ground truth data. Consistent with methods reported in the literature, we used Spearman’s rank correlation coefficient to compare a predicted feature vector 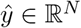 with the ground truth *y* ϵ ℝ^*N*^, transformed to ranks 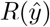 and *R*(*y*):^17,19,20,22^

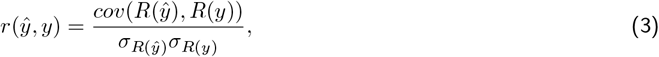

where 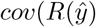, *R*(*y*)) is the covariance of the rank variables, and 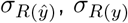 are the standard deviations.

To compute scores across the 10 test partitions, we first computed the mean Spearman’s rank correlation coefficient per individual feature. We then reported the average for the top predicted features. The error was calculated as the mean standard deviation across features.

### 2.4. Training and evaluation of inflammatory bowel disease classifiers

To evaluate the applicability of meta-omics prediction models, we used the predicted features for the downstream task of inflammatory bowel disease (IBD) prediction. All classification tasks were performed using scikit-learn’s RandomForestClassifier (v1.4.1.post1), trained using random search cross-validation (Supplementary Table S6) with 50 iterations and a random state equal to the seed corresponding to each train/test partition (all random seeds are listed in Supplementary Table S3). We used 5 stratified cross-validation folds, divided based on study participants.

For each paired dataset, we reported the accuracy of IBD classifiers trained on the predicted data to that of classifiers trained on the input data used to generate the corresponding predictions. We additionally benchmarked these results against classifiers trained on ground-truth datasets of metatranscriptomics, metaproteomics, and metabolomics data. The training and test partitions were kept as the same ones used to train the cross-omics regression models (Section 2.3). To provide a fair comparison, for each meta-omics output type (mTx, mPx, mBx), we downsampled the classifier training sets to the size of the smallest paired meta-omics dataset. In addition, each train and test set was downsampled to equal class proportions (IBD and healthy control).

## 3. Results

### 3.1. A benchmarking pipeline for meta-omics prediction

To provide a standardized way to assess the performance of machine learning models to infer one meta-omics data type from another, we created a systematic evaluation protocol (see Figure 1).

**Figure 1:**
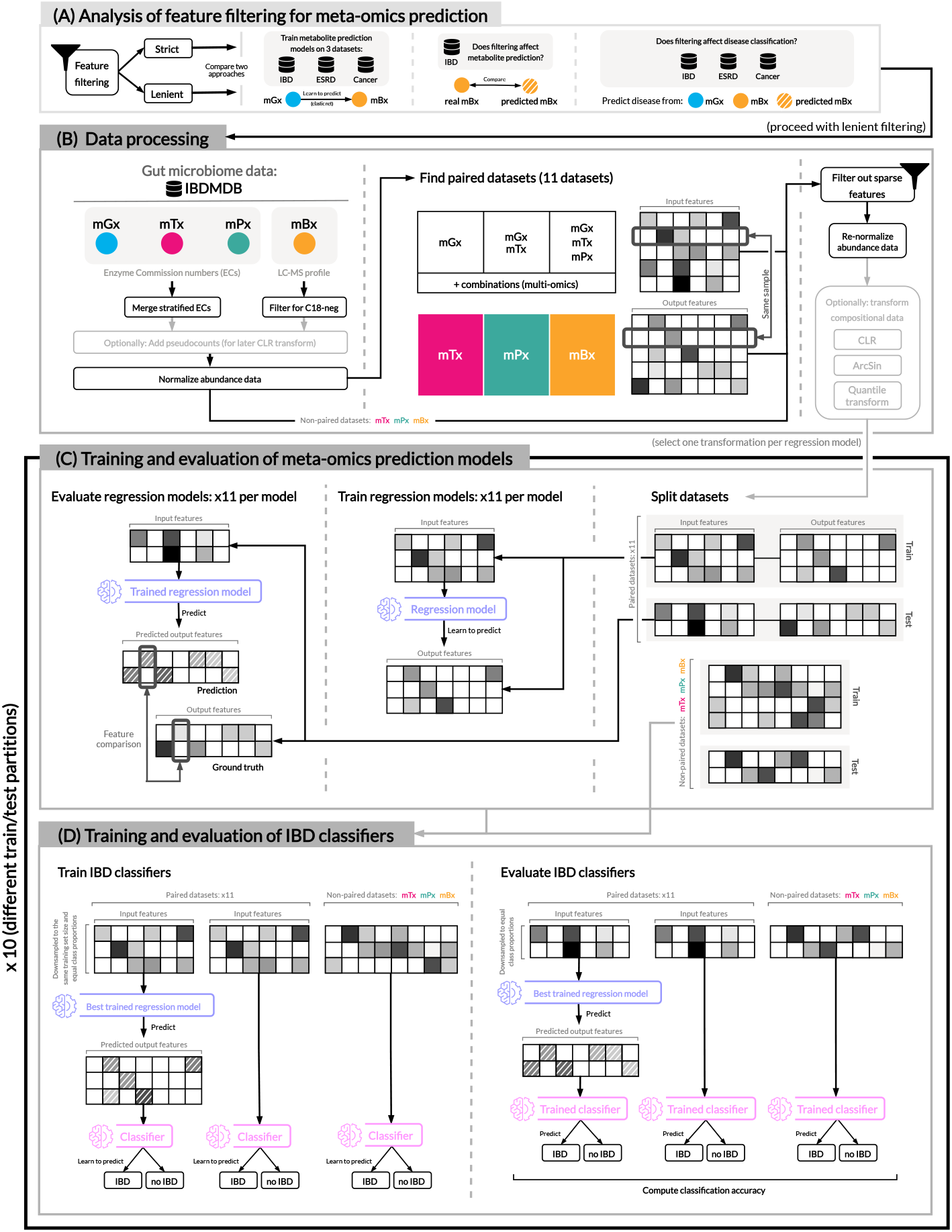
Overview of our experimental set-up. (A) We perform a pre-evaluation of MelonnPan [17] on three paired metagenomics and metabolomics datasets (Supplementary Table S1), comparing two filtering approaches for microbial features. In our main experimental pipeline, we use pre-processed gut microbiome data (B) to train regression models as meta-omics predictors (C), and subsequently evaluate these predictions for the downstream task of IBD prediction (D). Abbreviations: IBD (inflammatory bowel disease), ESRD (End-Stage Renal Disease), IBDMDB (The Inflammatory Bowel Disease Multi’omics Database), mGx (metagenomics), mTx (metatranscriptomics), mPx (metaproteomics), mBx (metabolomics), ECs (enzyme commission numbers), LC-MS (liquid chromatography-mass spectrometry), CLR (centered log-ratio).

First, we evaluated the effect of feature filtering on model performance (Figure 1A). We used an existing prediction method, i.e. MelonnPan, proposed by Mallick et al.^17^, to predict metabolite profiles from metagenomics data, using three datasets focused on different microbiome-associated conditions: inflammatory bowel disease, end-stage renal disease and colorectal cancer.^13,23,24^ We compared two filtering approaches for sparse features: strict filtering, which resulted in a high number of microbial features being filtered out, and lenient filtering, in which fewer features were filtered out. For each filtering procedure, experimental and predicted data were also used to classify disease phenotypes.

Although the quality of metabolite predictions was invariant to the filtering approach (Supplementary Figure S1A), we found that the filtering procedure did have an impact on downstream classification performance (Supplementary Figure S1B and C). Across all three datasets, we noticed a significant drop in performance for classifiers trained on experimental metabolomics data, with the strict filtering approach. This suggests that features important for distinguishing phenotype were discarded as a result of strict filtering, so we reported results using the lenient filtering approach for the remainder of the manuscript.

Next, we designed an experimental pipeline to assess the utility of machine learning models for meta-omics prediction, integrating three main components: processing of paired microbiome data (Figure 1B), training and evaluation of meta-omics prediction models (Figure 1C), and, lastly, classification of inflammatory bowel disease with predicted data (Figure 1D). Using multi-omics data included in the Inflammatory Bowel Disease Multi’omics Database (IBDMDB), we selected paired samples from several combinations of meta-omics modalities, resulting in 11 paired datasets that enabled the prediction of transcript, protein and metabolite abundances from various input types. This data was then filtered for sparsity and transformed according to standard procedures for compositional data (see Methods). Afterwards, the following procedure was repeated ten times, to ensure that the reported performance was robust. Each processed dataset was divided into a training and test set, selected based on participant IDs, since the presence of samples from the same patient in both the training and the test set might have resulted in overfitting. Five multi-output regression models were then trained and evaluated for the task of meta-omics feature prediction: an elastic net^17^, a feed-forward network^19^, a sparse encoder-decoder^18^, a deep neural network and a random forest regression baseline. The best model, i.e. the elastic net, was further used to generate meta-omics predictions in the last step in our pipeline, in which we compared the performance of IBD classifiers trained on three types of input data: predicted meta-omics data, the data from which it was predicted, and the ground-truth.

### 3.2. Machine learning models can reliably predict a subset of meta-omics features

To assess the generalization performance of machine learning pipelines designed for metabolite prediction, we first trained and evaluated several models from the literature on the task of predicting transcript and protein abundances, in addition to metabolite abundances. Average scores across 10 different train/test partitions were computed for 6 single-omics input-output combinations and 3 models from the literature MelonnPan, MiMeNet, and SparseNED.^17–19^ To these we add two new classifiers: (i) a deep neural network (Deep NN, as described in Supplementary Note A.1) and (ii) a random forest regressor (Figure 2) as an additional baseline model. Following a pattern established in the literature, we plotted the performance of cross-omics regression models for the 50 best predicted features (Figure 2A).^17,19,20,22^

**Figure 2:**
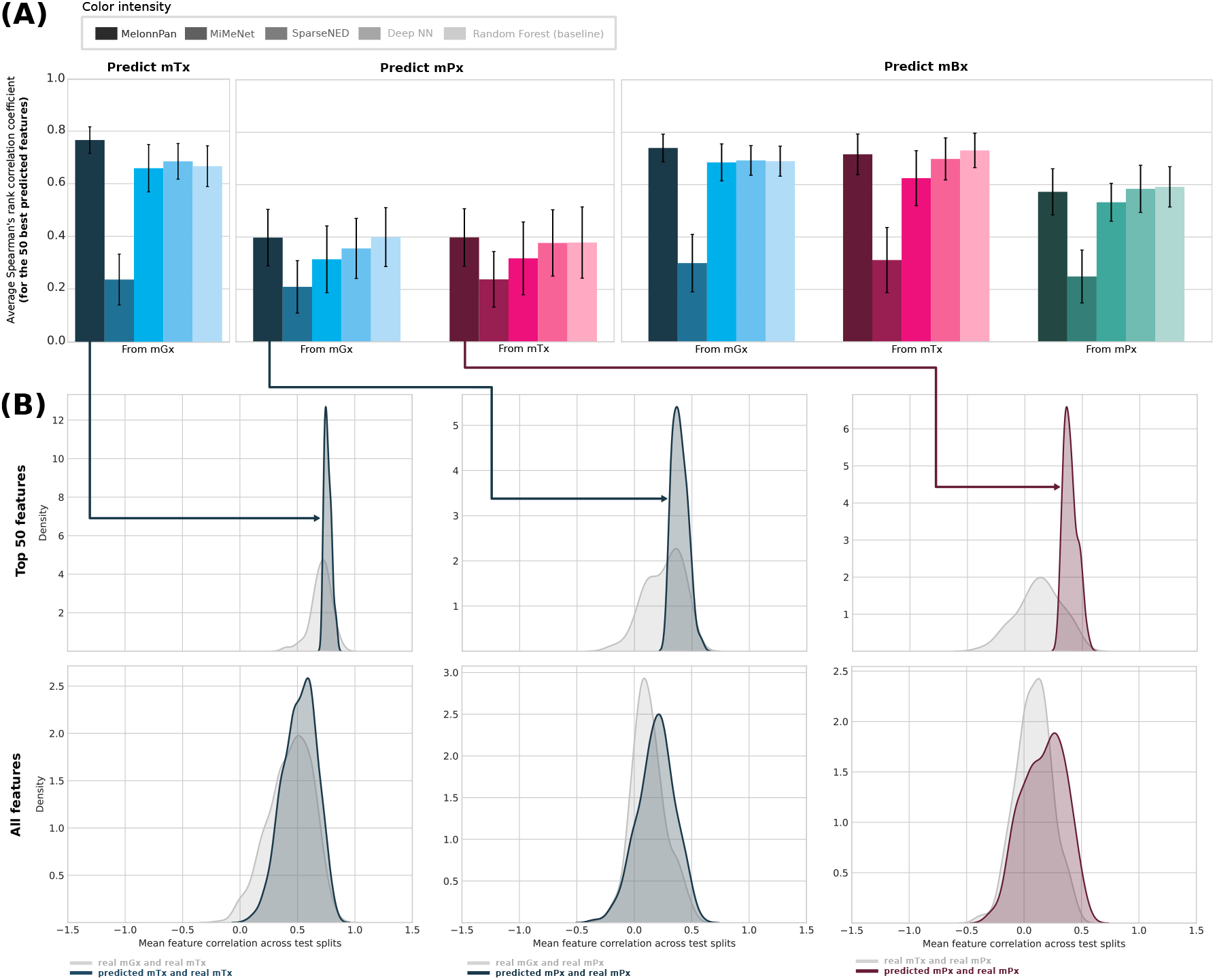
**(A)** Mean test performance results of cross-omics regression models on several prediction tasks, calculated across 10 different dataset partitions. The average Spearman’s rank correlation coefficient was calculated for the 50 best predicted features for each output type. **(B)** For metatranscriptomics (mTx) and metaproteomics (mPx) predictions generated by MelonnPan [17], we also plot kernel density estimates comparing correlations between the input data and the ground-truth mTx/mPx data with those computed between the predicted data and the ground truth data. We perform this analysis on the 50 best predicted features for each output type, as well as all predicted features. Input types are represented through different colors, while cross-omics models are represented using different color intensities. Abbreviations: neural network (NN), metagenomics (mGx), metatranscriptomics (mTx), metaproteomics (mPx), metabolomics (mBx).

Considering the top predictions, cross-omics regression models performed similarly in predicting metatranscriptomics and metabolomics (Figure 2A), with elastic nets achieving the highest average correlations, measuring 0.77 and 0.74, respectively. Protein abundances (mPx) were the most challenging to predict, with an average correlation coefficient of at most 0.4. In general, architectures like elastic nets and random forests were more robust across input-output combinations and performed best among the benchmarked models.

To investigate whether machine learning models provide more accurate estimations of transcript and protein abundances compared to the “gene-to-transcript-to-protein” assumption, we generated density plots comparing the distributions of correlations between different types of meta-omics data (Figure 2B). Top predicted transcript and protein abundances were significantly more highly correlated with the ground-truth data, when compared to the distribution of correlations between the input data and the ground-truth. This shows that a subset of features (transcripts or proteins) can be more reliably predicted using machine learning approaches, rather than relying on the assumption that genes encoded in the metagenome will be transcribed into mRNA and subsequently translated into protein. When plotting correlations for all features, this was still the case, but to a much lesser extent, ultimately indicating that only a subset of features can be reliably predicted.

### 3.3. Multi-omics integration does not lead to better predictions

To determine whether using a combination of different types of input features leads to better predictions of protein and metabolite abundances, we additionally trained MelonnPan, the overall best performing model identified in the previous section, on multi-omics input. Figure 3 shows a comparison between single-omics and multi-omics input in predicting metaproteomics and metabolomics. Results for other input types, such as taxonomic profiles and pathways derived from metagenomics, are recorded in Supplementary Table S7.

**Figure 3:**
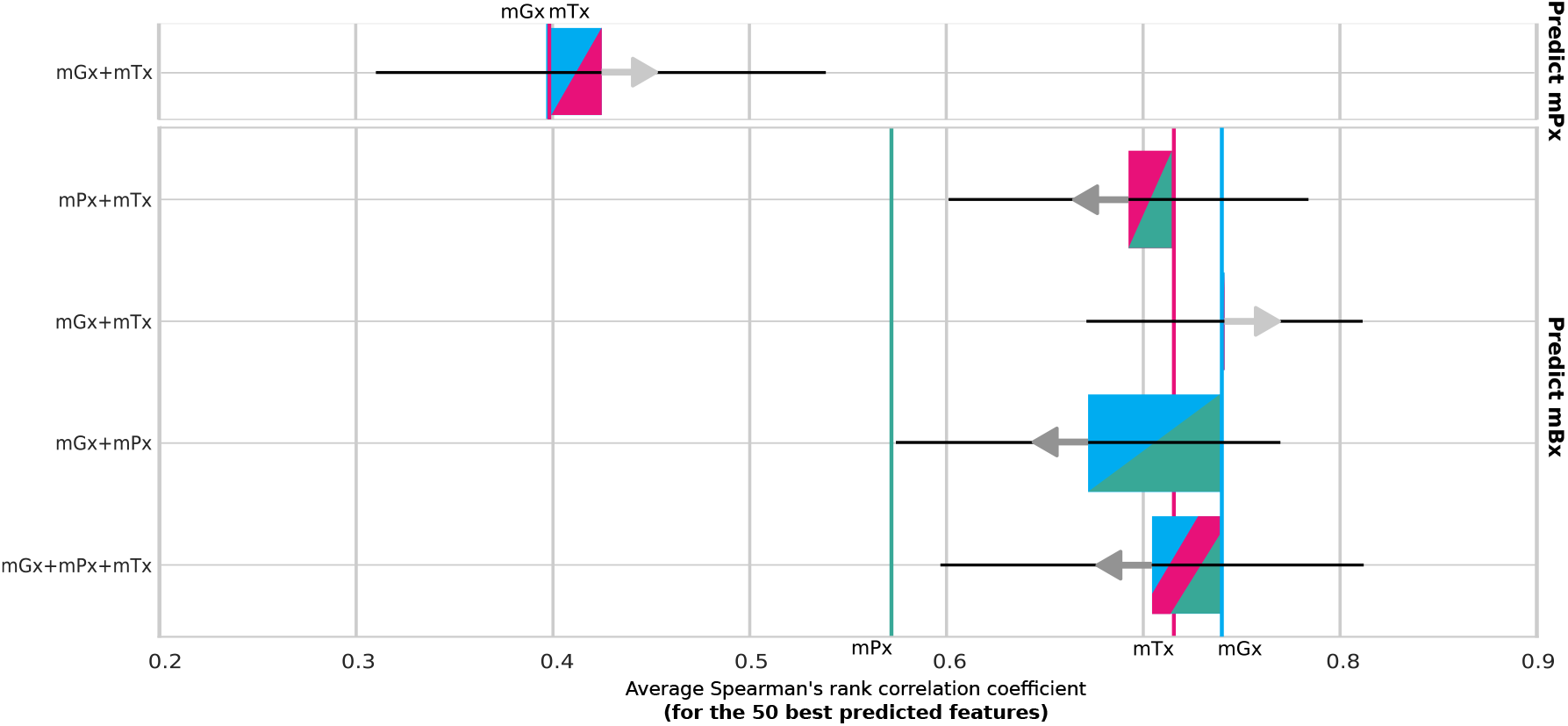
Performance comparison of multi-omics and single-omics input data using MelonnPan [17]. Results of single-omics input types are shown as vertical colored lines. Model performance on multi-omics data is indicated relatively to the best single-omics input type in a combination. The combination is displayed as a two- and three-color diagonally spliced bar with colors indicating including respective data types. Improvements or downgrades in performance are indicated with arrows and the size of the bar. Abbreviations: metagenomics (mGx, blue), metatranscriptomics (mTx, pink), metaproteomics (mPx, green), metabolomics (mBx, yellow).

While metaproteomics predictions marginally improved when combining metagenomics and metatranscriptomics, with 2% higher average correlation, combining single-omics modalities did not lead to more accurate predictions of metabolite abundances. Comparable performance was obtained when using metagenomics data processed in the form of pathways or species-level taxonomic profiles (Supplementary Table S7).

We also designed a more elaborate multi-omics integration scheme, using an auto-encoder trained with a joint reconstruction and regression loss (Supplementary Note A.2 and Supplementary Figure S4), but experiments did not show promising results. Therefore, we did not pursue this line of research further. Supplementary Table S8 shows the performance of MelonnPan trained on concatenated multi-omics, compared to the embeddings learned by the auto-encoder architecture. Although we observed a decline in prediction accuracy, the models trained on latent features were more robust, as suggested by the very low variation in model performance across test partitions.

### 3.4. Core set of well-predicted features is robust to input perturbations

We evaluated the robustness of cross-omics models through an analysis of well-predicted features across dataset partitions and input types (Figure 4(A), (B) and (C)). We limited this analysis to results produced by MelonnPan, as we found this model performed best overall for the task of cross-omics prediction (Section 3.2). However, additional experiments were performed with a deep neural network model (Supplementary Note A.1), to study the effect of feature selection on model performance (Figure 4(D)).

**Figure 4:**
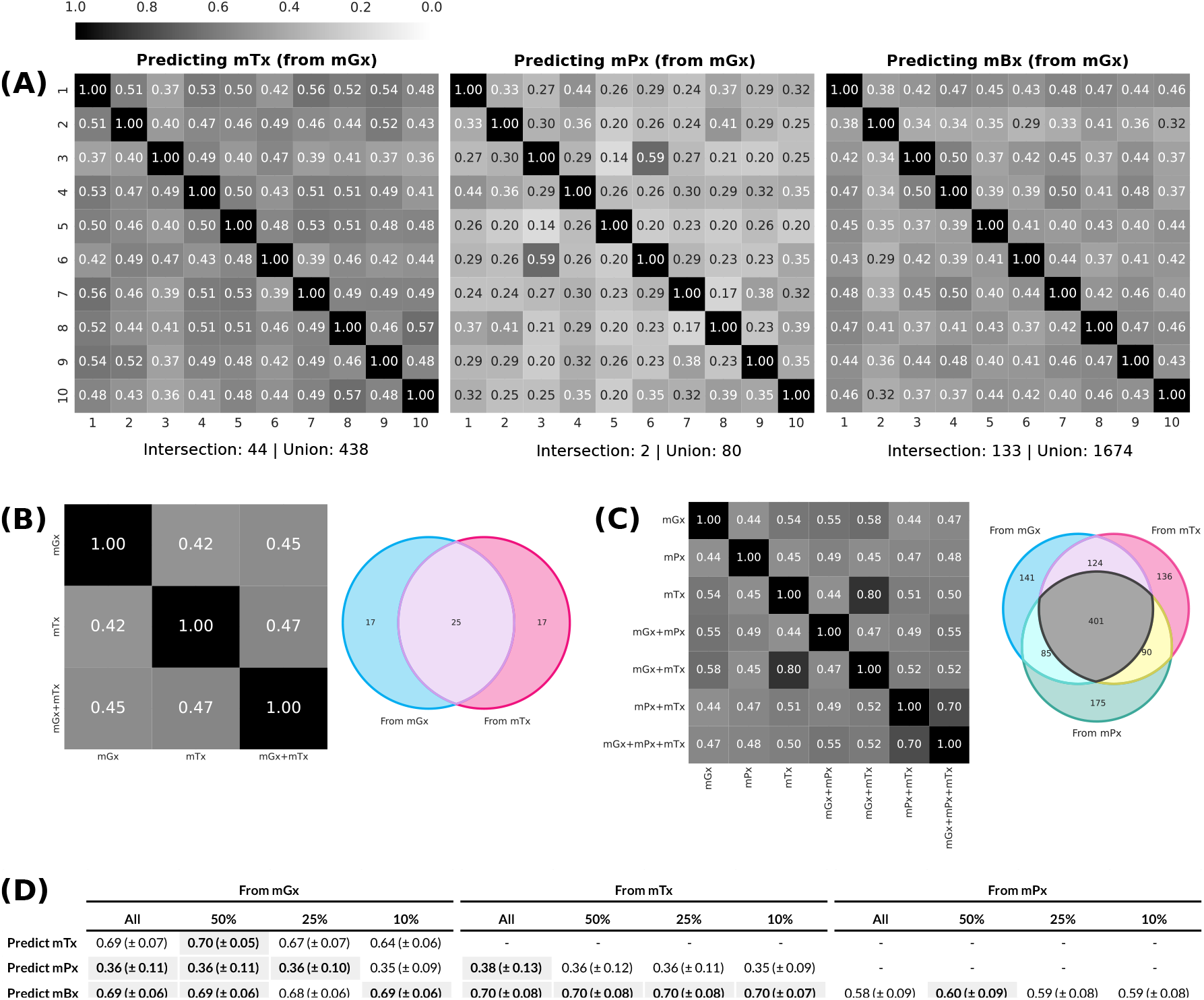
**(A)** Jaccard similarities between the sets of the 25% best predicted features by MelonnPan [17] for each output type (mTx, mPx, mBx), compared across 10 different train/test partitions. All predictions were generated from mGx data. **(B)** Jaccard similarities and Venn diagram of the sets of the 25% best predicted proteins, compared across input types. **(C)** Jaccard similarities and Venn diagram of the sets of the 25% best predicted metabolites, compared across input types. **(D)** Performance of a deep neural network model (Supplementary Section A.1) trained on different feature subsets (all, 50%, 25% and 10%), based on a pre-training step for feature selection (Supplementary Section A.3). The best results for each input-output combination are highlighted. Abbreviations: metagenomics (mGx), metatranscriptomics (mTx), metaproteomics (mPx), metabolomics (mBx).

A pairwise comparison of the top 25% well-predicted features across train/test partitions showed that these subsets share a selection of features, with some features being well-predicted across all test splits. While Figure 4(A) only shows predictions generated from mGx data, we found this to be the case for all single-omics input types (see Supplementary Figure S2). For each output type, a small set of features was found to be consistently well predicted across dataset partitions. For metaproteomics, this number was especially low (2.5% of the feature union). Glutamate dehydrogenase (1.4.1.3) and DNA-directed RNA polymerase (2.7.7.6) were the two enzymes included in this subset. Low abundance levels for these two enzymes have been associated with IBD diagnosis, with glutamate dehydrogenase being linked to *Clostridium difficile* infections in IBD patients.^29–31^

Some well-predicted features were also shared across single-omics and multi-omics input types (Figure 4(B) and (C)). In total, 25 proteins were well-predicted from both metagenomics and metatranscriptomics data, while 401 metabolites were well-predicted from metagenomics, metatranscriptomics and metaproteomics data. We also did not find significant correlations between feature variance and prediction quality (Supplementary Figure S3).

However, regardless of what makes features easy to predict, these results imply that there is a core subset of features, for each output type, that can be reliably predicted. Consequently, we hypothesized that training a model on just a subset of features would lead to better predictions, as the trade-off between data dimensionality and the number of samples would be more balanced in that case. Note that a multi-output elastic net such as Melonnpan would not be a suitable model to perform such an experiment. This is because this kind of architecture combines outputs from multiple independent single-output regression models, and, as such, reducing the number of output features would not have any effect on how accurately an individual feature can be predicted.^32^ On the other hand, a neural network architecture learns all output features simultaneously, so the number of output features matters during training.

To that end, we first ran a pre-training iteration which consisted of training 10 random forest models on different cross-validation splits, averaging feature correlations and retaining only a proportion of the top features (Supplementary Note A.3 and Supplementary Figure S5). We then trained a deep neural network (Supplementary Note A.1), restricting the output to subsets containing 50%, 25% and 10% of features based on individual correlations obtained during pre-training (Figure 4(D)). In addition to learning dependencies between output features, deep architectures were shown to be better at bypassing the curse of dimensionality, particularly when modeling compositional functions.^33^ Ultimately, network performance did not improve when training on a smaller set of output meta-omics features, but we also did not observe a decline in prediction accuracy (Figure 4D).

### 3.5. Predicted meta-omics data can classify phenotypes with performance comparable to experimental data

Lastly, to demonstrate an application of cross-omics regression models, we tested whether predictions generated by these models could be used for the downstream task of inflammatory bowel disease prediction. We compared the classification performance of random forest classifiers trained on input and predicted data, as well as experimental data of the same modality as the predictions (Figure 5). To ensure a fair comparison, datasets were downsampled to the same size and equal class proportions (IBD and healthy control).

**Figure 5:**
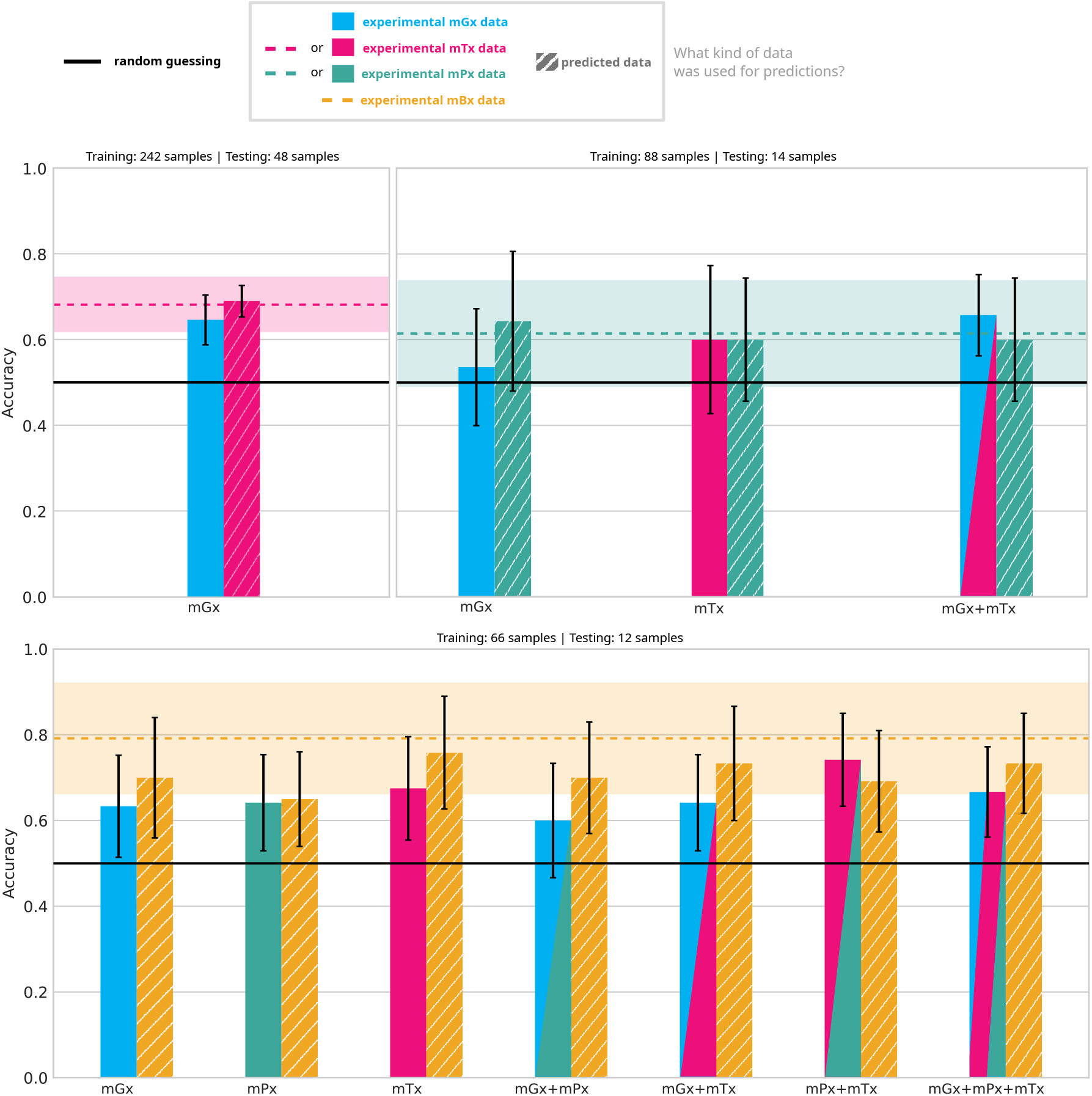
Accuracy of random forest classifiers on the binary task of inflammatory bowel disease prediction, averaged across 10 test partitions. Left bars indicate input types, while right, striped bars, indicate the output predicted from these inputs. Predicted data was generated with MelonnPan [17]. Abbreviations: metagenomics (mGx), metatranscriptomics (mTx), metaproteomics (mPx), metabolomics (mBx).

Overall, predicted features could be used to distinguish between IBD and the healthy control, with accuracies comparable to those obtained with experimental data. Metabolomics was identified as the best predictor of IBD, with a mean accuracy of almost 80%. Using metabolite abundances predicted from metatranscriptomics resulted in classifiers closely matching this performance. Similarly, classifiers trained with transcript and protein abundances generated from metagenomics data also led to improved performance.

In the majority of cases, predicted features were better predictors of inflammatory bowel disease compared to the input data from which they were generated. Using input data as opposed to predicted data was only preferred in multiomics settings. For instance, a combination of metagenomics and metatranscriptomics was preferable to predicted and experimental metaproteomics data and classifying inflammatory bowel disease samples. However, it was not a general trend that using more features, from diverse meta-omics modalities, led to better classification performance. A combination of metagenomics, metatranscriptomics and metaproteomics data resulted in less accurate classification compared to predicted metabolomics data.

## 4. Discussion

Our results have shown that metagenome-to-metabolite models can be generalized to meta-omics types. We found that regression models for cross-omics prediction are able to accurately predict a subset of features, whether those features are transcripts, proteins or metabolites. Although metaproteomics prediction was challenging, our experiments showed that metatranscriptomics and metabolomics features were reliably predicted. This is expected, given the fact that metaproteomics was characterized by the highest sparsity and that paired data available to train models that predict metaproteomics was most scarce. Good performance for metatranscriptomics prediction is also not entirely surprising, given the similarities between the measurement techniques for metagenomics and metatranscriptomics, i.e. next-generation sequencing technologies. Ultimately, we were able to validate similar results from the literature on metabolomics prediction.^17–20,22^

Our experiments also confirmed that machine learning models can provide reliable insights into metatranscriptomics and metaproteomics. Feature correlations between ground-truth and predicted data were generally higher than those obtained between genes and transcripts, or transcripts and proteins. Notably, this effect was less pronounced when all features were considered, as opposed to just the well-predicted ones. One explanation for this result is that machine learning becomes challenging when the number of features is high relative to the number of samples. This is particularly an issue with microbiome data, which is difficult to collect, sparse and high-dimensional, resulting in datasets with few samples and many features.^34–36^

We additionally evaluated these prediction models when trained on multi-meta-omics input, and generally observed a decrease in performance. We note that the number of samples available for training decreased with the amount of meta-omics modalities involved, and that likely also had an influence on these results. We expect that further investigation into multi-omics integration in a meta-omics setting should lead to more reliable predictors, given recent success in deep learning multi-omics integration for single-omics.^37–39^

As a potential solution to bypass the curse of dimensionality for this multi-output regression problem, we set up experiments to train a neural network using subsets of output features, based on how well they could be predicted in a preliminary cross-validation round. Unlike standard multi-output regression models, which essentially stack together single-output regressors, a neural network should be able to exploit dependencies between microbial features in the output layer.^32^ Although our results did not fully support the hypothesis that a smaller output dimensionality would lead to more streamlined model training, we found that performance also did not decrease with fewer features. This suggests that essential dependencies between predicted features were preserved during the feature selection process.

More importantly, our results suggest that a core subset of output features (transcripts, proteins and metabolites) can be predicted reliably regardless of training set composition and meta-omics input types. Analyzing such features independently may benefit researchers who wish to gain aspects into other meta-omics modalities, in cases when only metagenomics data is available. However, the characterization of well-predicted features remains a largely unanswered question, requiring more in-depth analysis, from a biological and statistical point of view.

Finally, we proposed that predicted meta-omics features could be used to distinguish between different phenotypes. For the binary task of IBD classification, this hypothesis was confirmed. Some predicted datasets lead to classification performance similar to that obtained using experimental data, and generally better than that obtained using the input data at the basis of those predictions.

Classifying IBD and healthy samples on this dataset was a challenging task, particularly due to the limited number of training samples. This is a result of downsampling to equal class proportions, on top of downsampling to the size of the smallest paired dataset for each output type. Additionally, the IBD samples in IBDMDB were not all collected from patients with active disease, making it harder to distinguish between the two classes of samples.^26^ Our initial experiments on other paired metagenomics and metabolomics datasets (Section 3.1) also provided evidence that this issue is partially dataset-related, as random forest classifiers with no hyperparameter tuning were able to achieve good performance with experimental metabolomics data, even for more challenging classification tasks. Ultimately, in a clinical, applied setting, inflammatory bowel disease is generally not a straightforward diagnosis.^40^ Our aim is not for this classification performance to be competitive with the state-of-the-art in microbiome-based disease prediction, which relies on more complex, deep model architectures (see, for instance, Liao et al.^41^ and Shi et al.^42^), but that it serves as a proof-of-concept for the utility of predicted meta-omics data in microbiome research.

## Data availability statement

This study uses data available as part of the The Inflammatory Bowel Disease Multi’omics Database^43^ and The Curated Gut Microbiome Metabolome Data Resource (v2.1.0)^44^.

