## Supplemental for "Predicted meta-omics: a potential solution to multi-omics data scarcity in microbiome studies"

### A. Supplementary methods

#### A.1. Deep fully-connected neural network

We trained 36 deep neural network (Deep NN) architectures, with various numbers of hidden layers, different data augmentation factors and loss functions. In the end, the architecture that was used throughout our experiments included 3 fully connected hidden layers and a loss function including equal proportions of Pearson's correlation and the mean-squared error (MSE). Additionally, training was performed without any data augmentation. For a comprehensive benchmark of different architectures and augmentation factors, see Supplementary Table S9.

##### A.1.1. Data augmentation

We augmented the paired cross-omics datasets using an approach inspired by the Aitchison mixup described by Gordon-Rodriguez *et al.* [1]. However, one important distinction is that the authors describe augmentation of compositional data in the simplex, before transformations are applied, but we apply augmentation on the transformed data. Data was transformed using the quantile transformation, as described in Section 2.2.

Let $x_i$ and $x_j \in \mathbb{R}^D$ be two input samples, with corresponding output samples $y_i$ and $y_j \in \mathbb{R}^M$. We construct augmented data points $x'$ and $y'$ using a linear combination:

$$x' = \lambda \cdot x_i + (1 - \lambda) \cdot x_j,$$
$$y' = \lambda \cdot y_i + (1 - \lambda) \cdot y_j,$$

where $\lambda \in [0, 1]$ and $i, j \in \{1, ..., N\}$. To generate multiple data points, $i$ and $j$ are chosen randomly, and $\lambda$ is sampled from a uniform distribution.

##### A.1.2. Architecture

Layer dimensions were chosen based on input size. To that end, we constructed architectures of the form: `input_size` – $\lfloor$`1.25` · `input_size`$\rfloor$ – ... – $\lfloor$`1.25` · `input_size`$\rfloor$ – $\lfloor$`2.5` · `input_size`$\rfloor$ – `output_size`. Layer norm and ReLU were applied after each layer, excluding the output layer.

##### A.1.3. Loss function

We defined a loss function based on a combination of Pearson's correlation and the mean squared error, between the ground truth $Y \in \mathbb{R}^{N \times M}$ and the prediction $\hat{Y} \in \mathbb{R}^{N \times M}$:

$$L(Y, \hat{Y}) = \alpha_{MSE} \cdot MSE(Y, \hat{Y}) + \alpha_{corr} \cdot (1 - \rho(Y, \hat{Y})), \tag{4}$$

where $\rho(Y, \hat{Y})$ represents the average Pearson correlation coefficient between ground-truth and predicted features, $MSE(Y, \hat{Y})$ is the mean squared error between the ground truth and the prediction, and $\alpha_{MSE}$ and $\alpha_{corr} \in [0, 1]$.

To compute the mean squared error, we used `torch`'s (2.1.2.post300) MSE loss, from the `nn.functional` module, with "mean" reduction. To determine Pearson's correlation coefficient, we applied the CosineEmbeddingLoss from the same module. Prior to this, each feature was centered around its mean, and the data batch was transposed.

##### A.1.4. Training procedure

All network models (Supplementary Table S9) were constructed using the Pytorch Lightning API, version 2.2.1, with a random seed equal to 42. Training and validation sets were split based on study participants, as described in Section 2, using a shuffled split, with a random seed equal to 42. We used a batch size of 16 and the Adam optimizer, with a learning rate equal to `1e-4`, a patience of 3 for early stopping, and a maximum of 35 epochs.

#### A.2. Multi-omics autoencoder

As an alternative to naive feature concatenation, we trained an autoencoder model for multi-omics integration. A diagram of the architecture and loss function is included in Supplementary Figure S4. After training the network, we used the latent features to train the best-performing model in our benchmark, MelonnPan[2]. Below we provide details on the network architecture and the loss function used during training.

##### A.2.1. Architecture

As shown in Supplementary Figure S4, we divided the model architecture into two main parts: the autoencoder and a multi-layer perceptron, which takes as input the latent features of the autoencoder, and then predicts a meta-omics output. The autoencoder was organized using a symmetric architecture, with a hidden layer of dimension $\lfloor 0.75 \cdot \texttt{input\_size} \rfloor$, and a latent space of dimension $\lfloor 0.5 \cdot \texttt{input\_size} \rfloor$. The multi-layer perceptron included one hidden layer, of dimension $\lfloor 0.25 \cdot \texttt{input\_size} \rfloor$. Following a similar approach as described previously (Supplementary Section A.1.3), we applied layer norm and a ReLU activation after each layer, excluding the output layers.

##### A.2.2. Loss function

We constructed an architecture to enable learning of embeddings for the task of cross-omics prediction. To that end, we trained an autoencoder with a combined loss, integrating the reconstruction loss with a regression loss. This was inspired by the approach described by Hira *et al.* [3], who jointly trained a variational autoencoder and classifier for ovarian cancer, using a combined loss.

Let $X \in \mathbb{R}^{N \times D}$ be the input multi-meta-omics feature matrix, and let $Y \in \mathbb{R}^{N \times M}$ be the output meta-omics feature matrix. In addition, let $\hat{X} \in \mathbb{R}^{N \times D}$ be the prediction produced by the autoencoder, and let $\hat{Y} \in \mathbb{R}^{N \times M}$ be the prediction produced by the multi-layer perceptron. We computed the following loss:

$$L(X, \hat{X}, Y, \hat{Y}) = L(X, \hat{X}) + L(Y, \hat{Y}), \tag{5}$$

where $L(X, \hat{X})$ and $L(Y, \hat{Y})$ are defined as in equation 4.

##### A.2.3. Training procedure

We followed the same procedure as described in Supplementary Section A.1.4, with the exception that the maximum number of epochs was set to 50.

#### A.3. Feature selection

We designed a pre-training step to select a small set features to be used later during model training (Supplementary Figure S5). To that end, we split each training set into 10 training/validation partitions, and trained a random forest regressors on each partition. Feature correlations were subsequently calculated on the validation set, and each feature was assigned a score, equal to the mean across validation sets. Based on these scores, we retained a fraction of features to be later used for cross-omics model training.

### B. Commands for model training

```
# MelonnPan
# Training
Rscript train_metabolites.R -i [path_to_input_data] -g [path_to_output_data] -p 20 -o [path_to_output_folder]
# Making predictions with the re-trained model
Rscript predict_metabolites.R -w [path_to_trained_weights] -i [path_to_input_test_data] -o [path_to_output_folder]


# SparseNED
```

```
python3 main_cv_1dir.py --model BiomeAESnip --sparse 0.06 --learning_rate 0.01 --batch_size 20
     --latent_size 70 --activation "tanh_tanh" --data_type [dataset_name] --data_root [
    path_to_data_folder] --nonneg_weight --normalize_input

# MiMeNet
python3 MiMeNet_train.py -micro [path_to_input_data] -metab [path_to_output_data] -micro_norm
    None -metab_norm None -net_params None -external_micro + [path_to_input_test_data] -
    external_metab [path_to_output_test_data] -num_background 10 -num_run 5 -num_cv 5
```

**Listing 1:** Commands used to run "metagenomics-to-metabolomics" tools: MelonnPan [2], SparseNED[4] and MiMeNet [5].

### C. Supplementary tables

**Table S1:** Description of paired metagenomics and metabolomics datasets used for experimental validation and reproducibility testing.

| | | | | Lenient filtering | | Strict Filtering | | No filtering | |
|---|---|---|---|---|---|---|---|---|---|
| **Dataset name** | **Study** | **# train samples** | **# test samples** | **Input dim.** | **Output dim.** | **Input dim.** | **Output dim.** | **Input dim.** | **Output dim.** |
| CD, UC and HC | Franzosa et al. [12] | 102 | 60 | 1231 | 4541 | 722 | 1415 | 2113 | 8848 |
| ESRD and HC | Wang et al. [42] | 106 | 28 | 1401 | 274 | 795 | 94 | 56961 | 276 |
| Cancer and HC | Yachida et al. [43] | 202 | 52 | 1462 | 269 | 698 | 76 | 57702 | 450 |

**Table S2:** Datasets downloaded from the Inflammatory Bowel Disease Multi'omics Database (IBDMB), used for training and testing machine learning models in our experiments.

| Data description | Download link | Download date |
|---|---|---|
| Metaproteomics ECs | https://www.ibdmdb.org/downloads/products/HMP2/MPX/2017-03-20/HMP2_proteomics_ecs.tsv.gz | 16.11.2023 |
| Metagenomics ECs | https://www.ibdmdb.org/downloads/products/HMP2/MGX/2018-05-04/ecs_relab.tsv.gz | 16.11.2023 |
| Metabolomics | https://www.ibdmdb.org/downloads/products/HMP2/MBX/HMP2_metabolomics.biom | 16.11.2023 |
| Metatranscriptomics ECs | https://www.ibdmdb.org/downloads/products/HMP2/MTX/2017-12-14/ecs_relab.tsv.gz | 16.11.2023 |
| Taxonomic profile | https://www.ibdmdb.org/downloads/products/HMP2/MGX/2018-05-04/taxonomic_profiles.tsv.gz | 16.11.2023 |
| Patient metadata | https://ibdmdb.org/downloads/metadata/hmp2_metadata_2018-08-20.csv | 17.01.2024 |

**Table S3:** Metadata for the paired-omics datasets used in our experiments. Input-output pairs used for the results of the main manuscript are highlighted. Results for the other datasets are only reported as part of the supplementary material.

| | | | Input/output dim. after filtering | | | | | | Train/test # samples per seed | | | | | | | | | | | | | | | | | | | |
|---|---|---|---|---|---|---|---|---|---|---|---|---|---|---|---|---|---|---|---|---|---|---|---|---|---|---|---|---|
| Output | Input | # samples | Non-imputed data | | Imputed data | | Train/test mean # samples | | 2 | | 3 | | 5 | | 7 | | 11 | | 13 | | 17 | | 23 | | 29 | | 31 | |
| **mTx** | **mGx** | **733** | **983** | **851** | **992** | **851** | **584,5** | **148,5** | **577** | **156** | **587** | **146** | **619** | **114** | **600** | **133** | **569** | **164** | **589** | **144** | **570** | **163** | **582** | **151** | **597** | **136** | **555** | **178** |
| mTx | mGx_taxa | 731 | 130 | 850 | 130 | 850 | 582,8 | 148,2 | 575 | 156 | 586 | 145 | 618 | 113 | 599 | 132 | 567 | 164 | 587 | 144 | 568 | 163 | 580 | 151 | 595 | 136 | 553 | 178 |
| mTx | mGx_pa | 732 | 323 | 850 | 323 | 851 | 583,6 | 148,4 | 576 | 156 | 587 | 145 | 618 | 114 | 599 | 133 | 568 | 164 | 588 | 144 | 569 | 163 | 581 | 151 | 596 | 136 | 554 | 178 |
| mTx | mGx+mGx_taxa | 731 | 1113 | 850 | 1117 | 850 | 582,8 | 148,2 | 575 | 156 | 586 | 145 | 618 | 113 | 599 | 132 | 567 | 164 | 587 | 144 | 568 | 163 | 580 | 151 | 595 | 136 | 553 | 178 |
| mTx | mGx+mGx_pa | 732 | 1306 | 850 | 1315 | 851 | 583,6 | 148,4 | 576 | 156 | 587 | 145 | 618 | 114 | 599 | 133 | 568 | 164 | 588 | 144 | 569 | 163 | 581 | 151 | 596 | 136 | 554 | 178 |
| mTx | mGx_pa+mGx_taxa | 731 | 453 | 850 | 453 | 850 | 582,8 | 148,2 | 575 | 156 | 586 | 145 | 618 | 113 | 599 | 132 | 567 | 164 | 587 | 144 | 568 | 163 | 580 | 151 | 595 | 136 | 553 | 178 |
| **mPx** | **mGx** | **280** | **1000** | **281** | **1004** | **909** | **222,2** | **57,8** | **217** | **63** | **223** | **57** | **218** | **62** | **221** | **59** | **223** | **57** | **223** | **57** | **222** | **58** | **226** | **54** | **223** | **57** | **226** | **54** |
| mPx | mGx_taxa | 278 | 135 | 281 | 135 | 909 | 220,4 | 57,6 | 215 | 63 | 222 | 56 | 216 | 62 | 220 | 58 | 221 | 57 | 221 | 57 | 220 | 58 | 224 | 54 | 221 | 57 | 224 | 54 |
| mPx | mGx_taxa+mTx | 185 | 1009 | 274 | 1010 | 909 | 146,7 | 38,3 | 150 | 35 | 146 | 39 | 150 | 35 | 136 | 49 | 145 | 40 | 147 | 38 | 145 | 40 | 153 | 32 | 150 | 35 | 145 | 40 |
| **mPx** | **mTx** | **186** | **874** | **274** | **875** | **909** | **147,6** | **38,4** | **151** | **35** | **146** | **40** | **151** | **35** | **137** | **49** | **146** | **40** | **148** | **38** | **146** | **40** | **154** | **32** | **151** | **35** | **146** | **40** |
| mPx | mGx+mGx_pa | 278 | 1339 | 281 | 1329 | 909 | 220,4 | 57,6 | 215 | 63 | 222 | 56 | 216 | 62 | 220 | 58 | 221 | 57 | 221 | 57 | 220 | 58 | 224 | 54 | 221 | 57 | 224 | 54 |
| mPx | mGx_pa | 278 | 323 | 281 | 325 | 909 | 220,4 | 57,6 | 215 | 63 | 222 | 56 | 216 | 62 | 220 | 58 | 221 | 57 | 221 | 57 | 220 | 58 | 224 | 54 | 221 | 57 | 224 | 54 |
| mPx | mGx+mGx_taxa | 278 | 1151 | 281 | 1154 | 909 | 220,4 | 57,6 | 215 | 63 | 222 | 56 | 216 | 62 | 220 | 58 | 221 | 57 | 221 | 57 | 220 | 58 | 224 | 54 | 221 | 57 | 224 | 54 |
| **mPx** | **mGx+mTx** | **186** | **1902** | **274** | **1903** | **909** | **147,6** | **38,4** | **151** | **35** | **146** | **40** | **151** | **35** | **137** | **49** | **146** | **40** | **148** | **38** | **146** | **40** | **154** | **32** | **151** | **35** | **146** | **40** |
| mPx | mGx_pa+mGx_taxa | 278 | 458 | 281 | 458 | 909 | 220,4 | 57,6 | 215 | 63 | 222 | 56 | 216 | 62 | 220 | 58 | 221 | 57 | 221 | 57 | 220 | 58 | 224 | 54 | 221 | 57 | 224 | 54 |
| mPx | mGx_pa+mTx | 185 | 1194 | 274 | 1201 | 909 | 146,7 | 38,3 | 150 | 35 | 146 | 39 | 150 | 35 | 136 | 49 | 145 | 40 | 147 | 38 | 145 | 40 | 153 | 32 | 150 | 35 | 145 | 40 |
| **mBx** | **mGx** | **388** | **1038** | **3247** | **1040** | **3247** | **311,7** | **76,3** | **314** | **74** | **315** | **73** | **311** | **77** | **316** | **72** | **309** | **79** | **316** | **72** | **321** | **67** | **307** | **81** | **307** | **81** | **301** | **87** |
| **mBx** | **mGx+mPx** | **203** | **1277** | **3345** | **1912** | **3345** | **161,3** | **41,7** | **161** | **42** | **163** | **40** | **160** | **43** | **158** | **45** | **162** | **41** | **164** | **39** | **167** | **36** | **157** | **46** | **162** | **41** | **159** | **44** |
| **mBx** | **mGx+mPx+mTx** | **154** | **2196** | **3373** | **2830** | **3373** | **121,2** | **32,8** | **124** | **30** | **118** | **36** | **116** | **38** | **118** | **36** | **121** | **33** | **123** | **31** | **116** | **38** | **128** | **26** | **125** | **29** | **123** | **31** |
| mBx | mGx_taxa | 386 | 129 | 3240 | 129 | 3240 | 310,1 | 75,9 | 313 | 73 | 315 | 71 | 309 | 77 | 314 | 72 | 307 | 79 | 314 | 72 | 319 | 67 | 305 | 81 | 306 | 80 | 299 | 87 |
| mBx | mGx_taxa+mTx | 298 | 1002 | 3246 | 1002 | 3246 | 238,2 | 59,8 | 240 | 58 | 239 | 59 | 243 | 55 | 239 | 59 | 243 | 55 | 240 | 58 | 248 | 50 | 235 | 63 | 231 | 67 | 224 | 74 |
| **mBx** | **mTx** | **299** | **870** | **3252** | **871** | **3252** | **239,1** | **59,9** | **241** | **58** | **239** | **60** | **244** | **55** | **240** | **59** | **244** | **55** | **241** | **58** | **249** | **50** | **236** | **63** | **232** | **67** | **225** | **74** |
| mBx | mGx+mGx_pa | 386 | 1360 | 3240 | 1363 | 3247 | 310,1 | 75,9 | 313 | 73 | 315 | 71 | 309 | 77 | 314 | 72 | 307 | 79 | 314 | 72 | 319 | 67 | 305 | 81 | 306 | 80 | 299 | 87 |
| mBx | mGx_pa | 386 | 322 | 3240 | 323 | 3247 | 310,1 | 75,9 | 313 | 73 | 315 | 71 | 309 | 77 | 314 | 72 | 307 | 79 | 314 | 72 | 319 | 67 | 305 | 81 | 306 | 80 | 299 | 87 |
| mBx | mGx+mGx_taxa | 386 | 1167 | 3240 | 1169 | 3240 | 310,1 | 75,9 | 313 | 73 | 315 | 71 | 309 | 77 | 314 | 72 | 307 | 79 | 314 | 72 | 319 | 67 | 305 | 81 | 306 | 80 | 299 | 87 |
| **mBx** | **mPx+mTx** | **154** | **1158** | **3373** | **1792** | **3373** | **121,2** | **32,8** | **124** | **30** | **118** | **36** | **116** | **38** | **118** | **36** | **121** | **33** | **123** | **31** | **116** | **38** | **128** | **26** | **125** | **29** | **123** | **31** |
| **mBx** | **mGx+mTx** | **299** | **1922** | **3252** | **1925** | **3252** | **239,1** | **59,9** | **241** | **58** | **239** | **60** | **244** | **55** | **240** | **59** | **244** | **55** | **241** | **58** | **249** | **50** | **236** | **63** | **232** | **67** | **225** | **74** |
| mBx | mGx_pa+mPx | 202 | 602 | 3339 | 1233 | 3345 | 160,4 | 41,6 | 160 | 42 | 163 | 39 | 159 | 43 | 157 | 45 | 161 | 41 | 163 | 39 | 166 | 36 | 156 | 46 | 161 | 41 | 158 | 44 |
| mBx | mGx_pa+mGx_taxa | 386 | 451 | 3240 | 451 | 3240 | 310,1 | 75,9 | 313 | 73 | 315 | 71 | 309 | 77 | 314 | 72 | 307 | 79 | 314 | 72 | 319 | 67 | 305 | 81 | 306 | 80 | 299 | 87 |
| **mBx** | **mPx** | **307** | **282** | **3298** | **909** | **3298** | **242,4** | **64,6** | **242** | **65** | **241** | **66** | **249** | **58** | **241** | **66** | **239** | **68** | **247** | **60** | **238** | **69** | **243** | **64** | **242** | **65** | **242** | **65** |
| mBx | mGx_pa+mTx | 298 | 1190 | 3246 | 1194 | 3252 | 238,2 | 59,8 | 240 | 58 | 239 | 59 | 243 | 55 | 239 | 59 | 243 | 55 | 240 | 58 | 248 | 50 | 235 | 63 | 231 | 67 | 224 | 74 |
| mBx | mGx_taxa+mPx | 202 | 410 | 3339 | 1041 | 3339 | 160,4 | 41,6 | 160 | 42 | 163 | 39 | 159 | 43 | 157 | 45 | 161 | 41 | 163 | 39 | 166 | 36 | 156 | 46 | 161 | 41 | 158 | 44 |

**Table S4:** Metadata for the full meta-omics datasets used for IBD classification, before downsampling.

| | | Dimensionality after filtering | | | | Train/test # samples per seed | | | | | | | | | | | | | | | | | | | |
|---|---|---|---|---|---|---|---|---|---|---|---|---|---|---|---|---|---|---|---|---|---|---|---|---|---|
| **Data type** | **# samples** | **Non-imputed data** | **Imputed data** | **Train/test mean # samples** | | **2** | | **3** | | **5** | | **7** | | **11** | | **13** | | **17** | | **23** | | **29** | | **31** | |
| mTx | 735 | 851 | 851 | 586,1 | 148,9 | 578 | 157 | 589 | 146 | 620 | 115 | 602 | 133 | 570 | 165 | 590 | 145 | 572 | 163 | 584 | 151 | 599 | 136 | 557 | 178 |
| mPx | 449 | 283 | 909 | 358 | 91 | 350 | 99 | 356 | 93 | 359 | 90 | 360 | 89 | 356 | 93 | 358 | 91 | 361 | 88 | 364 | 85 | 353 | 96 | 363 | 86 |
| mBx | 546 | 3183 | 3183 | 433,5 | 112,5 | 439 | 107 | 430 | 116 | 434 | 112 | 433 | 113 | 438 | 108 | 430 | 116 | 435 | 111 | 430 | 116 | 434 | 112 | 432 | 114 |

**Table S5:** Comparison of data processing methods for three "metagenomics-to-metabolomics" models: MelonnPan[2], MiMeNet[5] and SparseNED[4]. We use the word "default" to refer to the data processing approach applied internally by the model, on normalized data. Some experiments were not performed. For example, MelonnPan already uses an arcsine and quantile transformation in its default pipeline, so we omitted that comparison. For each model, we highlight the data processing approach selected to report results for the model. Performance was measured using the average Spearman's rank correlation coefficient for the 50 best predicted features.

| | Predict mTx | Predict mPx | | Predict mBx | | |
|---|---|---|---|---|---|---|
| | From mGx | From mGx | From mTx | From mGx | From mPx | From mTx |
| **MelonnPan** | | | | | | |
| Normalized | 0.72 (± 0.07) | 0.35 (± 0.08) | 0.29 (± 0.06) | 0.64 (± 0.07) | 0.49 (± 0.06) | 0.61 (± 0.10) |
| ArcSin | - | - | - | - | - | - |
| CLR | 0.76 (± 0.05) | **0.42 (± 0.09)** | **0.47 (± 0.07)** | 0.72 (± 0.06) | **0.58 (± 0.08)** | **0.72 (± 0.08)** |
| Quantile transform | - | - | - | - | - | - |
| Model default | **0.77 (± 0.05)** | 0.40 (± 0.11) | 0.40 (± 0.11) | **0.74 (± 0.05)** | 0.57 (± 0.09) | **0.72 (± 0.08)** |
| **SparseNED** | | | | | | |
| Normalized | 0.53 (± 0.12) | 0.30 (± 0.12) | 0.26 (± 0.13) | 0.52 (± 0.13) | 0.40 (± 0.13) | 0.50 (± 0.13) |
| ArcSin | 0.64 (± 0.09) | **0.34 (± 0.12)** | 0.30 (± 0.13) | 0.65 (± 0.07) | 0.51 (± 0.09) | 0.59 (± 0.11) |
| CLR | 0.64 (± 0.10) | 0.32 (± 0.12) | 0.31 (± 0.12) | 0.54 (± 0.12) | 0.45 (± 0.07) | 0.61 (± 0.10) |
| Quantile transform | **0.66 (± 0.09)** | 0.31 (± 0.13) | **0.32 (± 0.14)** | **0.68 (± 0.07)** | **0.53 (± 0.07)** | **0.62 (± 0.11)** |
| Model default | - | - | - | - | - | - |
| **MiMeNet** | | | | | | |
| Normalized | **0.24 (± 0.11)** | 0.20 (± 0.09) | 0.23 (± 0.08) | 0.29 (± 0.09) | 0.27 (± 0.08) | 0.28 (± 0.09) |
| ArcSin | **0.24 (± 0.10)** | 0.21 (± 0.10) | 0.24 (± 0.11) | **0.30 (± 0.11)** | 0.25 (± 0.10) | **0.31 (± 0.12)** |
| CLR | - | - | - | - | - | - |
| Quantile transform | 0.23 (± 0.10) | 0.19 (± 0.10) | 0.22 (± 0.10) | 0.28 (± 0.10) | **0.32 (± 0.11)** | **0.31 (± 0.10)** |
| Model default | 0.23 (± 0.08) | **0.22 (± 0.12)** | **0.26 (± 0.10)** | 0.28 (± 0.09) | 0.31 (± 0.11) | 0.30 (± 0.11) |

**Table S6:** Grid values for hyperparameter tuning of random forest classifiers for IBD prediction.

| **Parameter name** | **Parameter values** |
|---|---|
| n_estimators | 64, 128, 256, 512, 1024 |
| max_depth | 16, 32, 64, 128, None |
| min_samples_split | 2, 4, 8 |
| min_samples_leaf | 1, 2, 4 |
| random_state | [equal to train/test partition seed] |

**Table S7:** Average Spearman's rank correlation coefficient of MelonnPan[2] predictions (top 50) for multiple single-omics and multi-omics input data types, including pathways (mGx_pa) and taxonomic profiles (mGx_taxa). The best results for each output type are highlighted.

| | Output type | | |
|---|---|---|---|
| **Input type** | **mTx** | **mPx** | **mBx** |
| mGx | 0.77 (± 0.05) | 0.40 (± 0.11) | 0.74 (± 0.05) |
| mGx+mGx_pa | 0.77 (± 0.05) | 0.41 (± 0.10) | 0.74 (± 0.05) |
| mGx+mGx_taxa | **0.78 (± 0.05)** | **0.42 (± 0.10)** | **0.75 (± 0.05)** |
| mGx+mPx | - | - | 0.67 (± 0.10) |
| mGx+mPx+mTx | - | - | 0.70 (± 0.11) |
| mGx+mTx | - | **0.42 (± 0.12)** | 0.74 (± 0.07) |
| mGx_pa | 0.72 (± 0.06) | 0.40 (± 0.10) | 0.71 (± 0.05) |
| mGx_pa+mGx_taxa | 0.76 (± 0.06) | 0.41 (± 0.11) | 0.74 (± 0.05) |
| mGx_pa+mPx | - | - | 0.66 (± 0.10) |
| mGx_pa+mTx | - | 0.41 (± 0.12) | 0.74 (± 0.07) |
| mGx_taxa | 0.75 (± 0.06) | 0.38 (± 0.11) | 0.72 (± 0.06) |
| mGx_taxa+mPx | - | - | 0.66 (± 0.09) |
| mGx_taxa+mTx | - | 0.41 (± 0.11) | **0.75 (± 0.07)** |
| mPx | - | - | 0.57 (± 0.09) |
| mPx+mTx | - | - | 0.69 (± 0.09) |
| mTx | - | 0.40 (± 0.11) | 0.72 (± 0.08) |

**Table S8:** Average Spearman's rank correlation coefficient of MelonnPan[2] predictions (top 50) for multi-omics input, comparing the model trained on a latent space, using the autoencoder in section A.2, to the model trained on naively concatenated multi-omics.

| Prediction task | MelonnPan trained on latent features | MelonnPan trained on concatenated multi-omics |
|---|---|---|
| mGx+mTx -> mPx | 0.16 (± 0.01) | 0.42 (± 0.12) |
| mGx+mPx -> mBx | 0.35 (± 0.01) | 0.67 (± 0.10) |
| mGx+mPx+mTx -> mBx | 0.40 (± 0.00) | 0.70 (± 0.11) |
| mGx+mTx -> mBx | 0.34 (± 0.01) | 0.74 (± 0.07) |
| mPx+mTx -> mBx | 0.40 (± 0.00) | 0.69 (± 0.09) |

**Table S9:** Spearman's rank correlation coefficient of the 50 best predicted features, for multiple combinations of network hyperparameters and input-output combinations, for a deep neural network model. Performance was computed on a validation set, separate from the test sets described in Section **??**. An augmentation factor equal to 1 indicates that no augmentation was applied, while an augmentation factor equal to $n > 1$ indicates that the final number of data points is equal to the initial size of the dataset multiplied by $n$. The best result for each input-output combination is highlighted.

| | **Predict mTx** | **Predict mPx** | | **Predict mBx** | | |
|---|---|---|---|---|---|---|
| **Model architecture** | **From mGx** | **From mGx** | **From mTx** | **From mGx** | **From mPx** | **From mTx** |
| Num. layers: 3 Aug. factor: 1 Loss: corr | 0.62 (± 0.07) | 0.32 (± 0.12) | 0.32 (± 0.15) | 0.61 (± 0.08) | 0.46 (± 0.11) | 0.59 (± 0.10) |
| Num. layers: 3 Aug. factor: 4 Loss: corr | 0.63 (± 0.07) | 0.30 (± 0.13) | 0.32 (± 0.14) | 0.61 (± 0.08) | 0.47 (± 0.12) | 0.59 (± 0.09) |
| Num. layers: 3 Aug. factor: 16 Loss: corr | 0.62 (± 0.07) | 0.32 (± 0.12) | 0.35 (± 0.14) | 0.59 (± 0.08) | 0.45 (± 0.13) | 0.56 (± 0.10) |
| Num. layers: 3 Aug. factor: 32 Loss: corr | 0.64 (± 0.07) | 0.32 (± 0.12) | 0.36 (± 0.14) | 0.61 (± 0.09) | 0.40 (± 0.11) | 0.59 (± 0.09) |
| Num. layers: 3 Aug. factor: 1 Loss: mse + corr | **0.67 (± 0.06)** | **0.40 (± 0.11)** | **0.43 (± 0.15)** | **0.68 (± 0.07)** | 0.53 (± 0.10) | 0.65 (± 0.06) |
| Num. layers: 3 Aug. factor: 4 Loss: mse + corr | **0.67 (± 0.06)** | 0.35 (± 0.12) | 0.40 (± 0.15) | 0.66 (± 0.07) | 0.50 (± 0.11) | 0.65 (± 0.07) |
| Num. layers: 3 Aug. factor: 16 Loss: mse + corr | 0.64 (± 0.06) | 0.32 (± 0.12) | 0.37 (± 0.14) | 0.62 (± 0.08) | 0.48 (± 0.10) | 0.60 (± 0.09) |
| Num. layers: 3 Aug. factor: 32 Loss: mse + corr | 0.62 (± 0.07) | 0.32 (± 0.13) | 0.37 (± 0.14) | 0.62 (± 0.07) | 0.44 (± 0.10) | 0.61 (± 0.09) |
| Num. layers: 3 Aug. factor: 1 Loss: mse | **0.67 (± 0.06)** | **0.40 (± 0.12)** | 0.42 (± 0.15) | **0.68 (± 0.07)** | 0.53 (± 0.10) | 0.67 (± 0.08) |
| Num. layers: 3 Aug. factor: 4 Loss: mse | **0.67 (± 0.06)** | 0.36 (± 0.12) | 0.40 (± 0.15) | 0.67 (± 0.07) | 0.53 (± 0.10) | 0.66 (± 0.07) |
| Num. layers: 3 Aug. factor: 16 Loss: mse | 0.64 (± 0.06) | 0.33 (± 0.12) | 0.37 (± 0.14) | 0.61 (± 0.08) | 0.48 (± 0.12) | 0.60 (± 0.08) |
| Num. layers: 3 Aug. factor: 32 Loss: mse | 0.63 (± 0.06) | 0.33 (± 0.12) | 0.37 (± 0.14) | 0.61 (± 0.09) | 0.44 (± 0.11) | 0.61 (± 0.09) |
| Num. layers: 4 Aug. factor: 1 Loss: corr | 0.62 (± 0.07) | 0.31 (± 0.12) | 0.31 (± 0.14) | 0.62 (± 0.08) | 0.44 (± 0.12) | 0.59 (± 0.09) |
| Num. layers: 4 Aug. factor: 4 Loss: corr | 0.64 (± 0.07) | 0.30 (± 0.12) | 0.33 (± 0.14) | 0.64 (± 0.08) | 0.49 (± 0.11) | 0.60 (± 0.08) |
| Num. layers: 4 Aug. factor: 16 Loss: corr | 0.63 (± 0.07) | 0.32 (± 0.12) | 0.35 (± 0.14) | 0.62 (± 0.08) | 0.42 (± 0.11) | 0.58 (± 0.09) |
| Num. layers: 4 Aug. factor: 32 Loss: corr | 0.66 (± 0.06) | 0.34 (± 0.12) | 0.36 (± 0.14) | 0.62 (± 0.08) | 0.42 (± 0.11) | 0.59 (± 0.09) |
| Num. layers: 4 Aug. factor: 1 Loss: mse + corr | 0.66 (± 0.07) | 0.36 (± 0.12) | 0.40 (± 0.15) | 0.65 (± 0.07) | 0.53 (± 0.12) | **0.70 (± 0.06)** |
| Num. layers: 4 Aug. factor: 4 Loss: mse + corr | **0.67 (± 0.06)** | 0.35 (± 0.11) | 0.39 (± 0.16) | 0.66 (± 0.07) | 0.50 (± 0.11) | 0.66 (± 0.07) |
| Num. layers: 4 Aug. factor: 16 Loss: mse + corr | 0.65 (± 0.06) | 0.34 (± 0.12) | 0.39 (± 0.14) | 0.63 (± 0.09) | 0.47 (± 0.12) | 0.60 (± 0.08) |
| Num. layers: 4 Aug. factor: 32 Loss: mse + corr | 0.65 (± 0.06) | 0.34 (± 0.12) | 0.36 (± 0.14) | 0.65 (± 0.07) | 0.44 (± 0.11) | 0.61 (± 0.08) |
| Num. layers: 4 Aug. factor: 1 Loss: mse | 0.66 (± 0.07) | 0.37 (± 0.12) | 0.39 (± 0.15) | 0.65 (± 0.09) | 0.53 (± 0.10) | 0.67 (± 0.07) |
| Num. layers: 4 Aug. factor: 4 Loss: mse | **0.67 (± 0.06)** | 0.37 (± 0.13) | 0.40 (± 0.14) | 0.67 (± 0.07) | 0.52 (± 0.12) | 0.67 (± 0.06) |
| Num. layers: 4 Aug. factor: 16 Loss: mse | 0.65 (± 0.06) | 0.35 (± 0.12) | 0.39 (± 0.14) | 0.63 (± 0.08) | 0.48 (± 0.11) | 0.62 (± 0.08) |
| Num. layers: 4 Aug. factor: 32 Loss: mse | 0.65 (± 0.06) | 0.35 (± 0.12) | 0.38 (± 0.14) | 0.64 (± 0.08) | 0.43 (± 0.11) | 0.63 (± 0.08) |
| Num. layers: 5 Aug. factor: 1 Loss: corr | 0.62 (± 0.08) | 0.29 (± 0.13) | 0.33 (± 0.15) | 0.64 (± 0.07) | 0.45 (± 0.12) | 0.59 (± 0.08) |
| Num. layers: 5 Aug. factor: 4 Loss: corr | 0.64 (± 0.07) | 0.30 (± 0.13) | 0.33 (± 0.15) | 0.63 (± 0.08) | 0.48 (± 0.12) | 0.61 (± 0.08) |
| Num. layers: 5 Aug. factor: 16 Loss: corr | 0.64 (± 0.07) | 0.32 (± 0.12) | 0.36 (± 0.14) | 0.62 (± 0.08) | 0.43 (± 0.11) | 0.60 (± 0.09) |
| Num. layers: 5 Aug. factor: 32 Loss: corr | **0.67 (± 0.06)** | 0.35 (± 0.13) | 0.35 (± 0.14) | 0.63 (± 0.09) | 0.42 (± 0.12) | 0.59 (± 0.09) |
| Num. layers: 5 Aug. factor: 1 Loss: mse + corr | 0.66 (± 0.07) | 0.36 (± 0.12) | 0.42 (± 0.15) | **0.68 (± 0.08)** | 0.53 (± 0.10) | 0.65 (± 0.08) |
| Num. layers: 5 Aug. factor: 4 Loss: mse + corr | **0.67 (± 0.06)** | 0.37 (± 0.13) | 0.40 (± 0.15) | 0.65 (± 0.07) | 0.51 (± 0.10) | 0.66 (± 0.06) |
| Num. layers: 5 Aug. factor: 16 Loss: mse + corr | 0.66 (± 0.06) | 0.34 (± 0.12) | 0.37 (± 0.14) | 0.63 (± 0.08) | 0.47 (± 0.11) | 0.63 (± 0.08) |
| Num. layers: 5 Aug. factor: 32 Loss: mse + corr | 0.66 (± 0.06) | 0.35 (± 0.12) | 0.38 (± 0.14) | 0.64 (± 0.08) | 0.45 (± 0.11) | 0.64 (± 0.08) |
| Num. layers: 5 Aug. factor: 1 Loss: mse | 0.66 (± 0.07) | 0.37 (± 0.12) | 0.41 (± 0.15) | 0.66 (± 0.07) | **0.55 (± 0.11)** | 0.68 (± 0.08) |
| Num. layers: 5 Aug. factor: 4 Loss: mse | **0.67 (± 0.06)** | 0.38 (± 0.11) | 0.41 (± 0.14) | 0.67 (± 0.07) | 0.52 (± 0.11) | 0.67 (± 0.07) |
| Num. layers: 5 Aug. factor: 16 Loss: mse | 0.66 (± 0.06) | 0.36 (± 0.12) | 0.39 (± 0.15) | 0.65 (± 0.08) | 0.48 (± 0.11) | 0.64 (± 0.07) |
| Num. layers: 5 Aug. factor: 32 Loss: mse | 0.66 (± 0.06) | 0.36 (± 0.12) | 0.38 (± 0.14) | 0.65 (± 0.07) | 0.45 (± 0.11) | 0.64 (± 0.07) |

### D. Supplementary figures

(A)

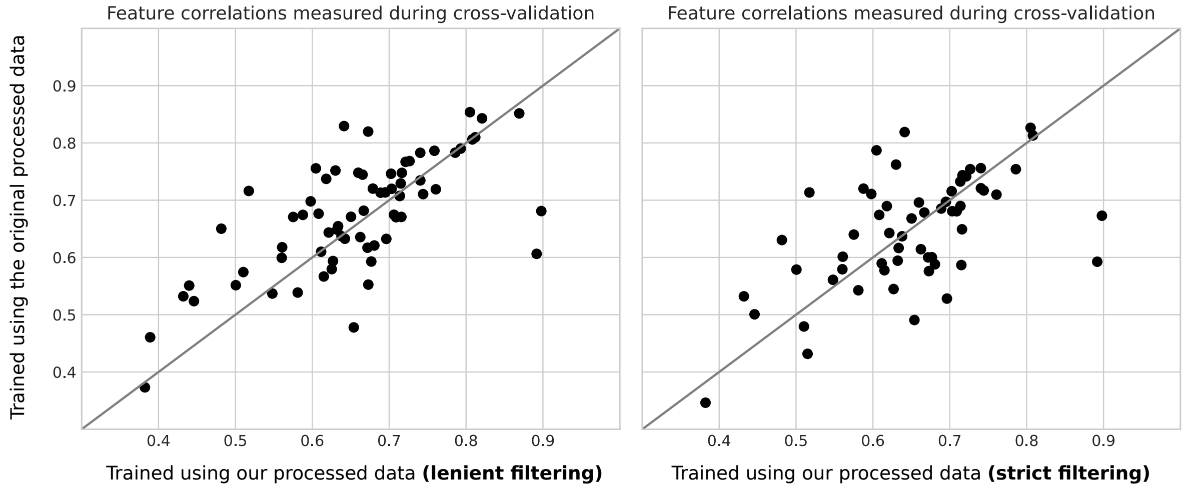


(B)

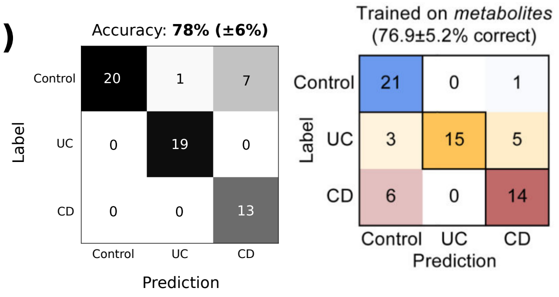


(C)

**Lenient filtering**

| Data | CD, UC and HC | ESRD and HC | Cancer and HC |
|---|---|---|---|
| mGx | 56% (±4%) | 93% (±3%) | 58% (±5%) |
| mBx | 78% (±6%) | 99% (±1%) | 67% (±3%) |
| Predicted mBx | 63% (±3%) | 93% (±6%) | 58% (±3%) |
| Predicted mBx (top features) | 62% (±2%) | 92% (±5%) | 57% (±3%) |

**Strict fltering**

| Data | CD, UC and HC | ESRD and HC | Cancer and HC |
|---|---|---|---|
| mGx | 58% (±4%) | 94% (±4%) | 54% (±4%) |
| mBx | 71% (±3%) | 86% (±5%) | 63% (±3%) |
| Predicted mBx | 58% (±5%) | 88% (±6%) | 57% (±5%) |
| Predicted mBx (top features) | 58% (±5%) | 88% (±6%) | 57 (±5%) |

**Figure S1: (A)** Spearman's rank correlations obtained by training MelonnPan [2] using our processed data (x-axis) and the data processed by the authors (y-axis). Correlations were computed during training across 10 folds of cross-validation. We include two feature filtering alternatives: less restrictive (left) and more restrictive (right). The original dataset was published by Franzosa *et al.* [6] (see also Supplementary Table S1). **(B)** On the left, a confusion matrix for the result highlighted in sub-figure (C). On the right, a confusion matrix taken from Figure 6 in the study published by Franzosa *et al.* [6]. **(C)** Performance of random forest classifiers for three different classification tasks, corresponding to the datasets [6–8] in Supplementary Table S1. Top mBx features were determined by MelonnPan during cross-validation, with a correlation cut-off equal to 0.3. Abbreviations: Crohn's disease (CD), ulcerative colitis (UC), healthy control (HC), end-stage renal disease (ESRD), metagenomics (mGx), metabolomics (mBx).

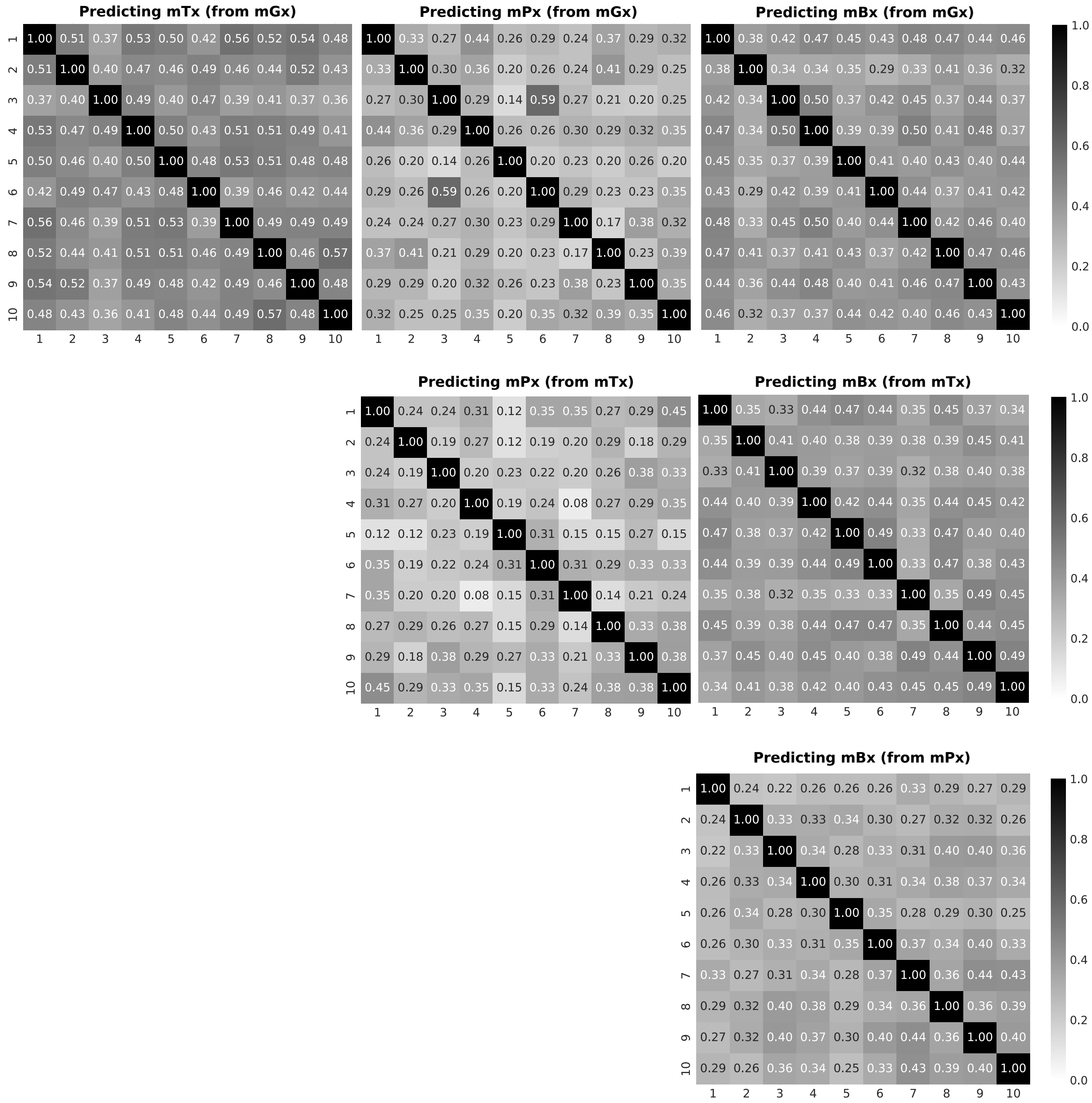


**Figure S2:** Jaccard similarities computed between sets of the top 25% well-predicted features across 10 train/test partitions. Predictions were generated with MelonnPan[2].

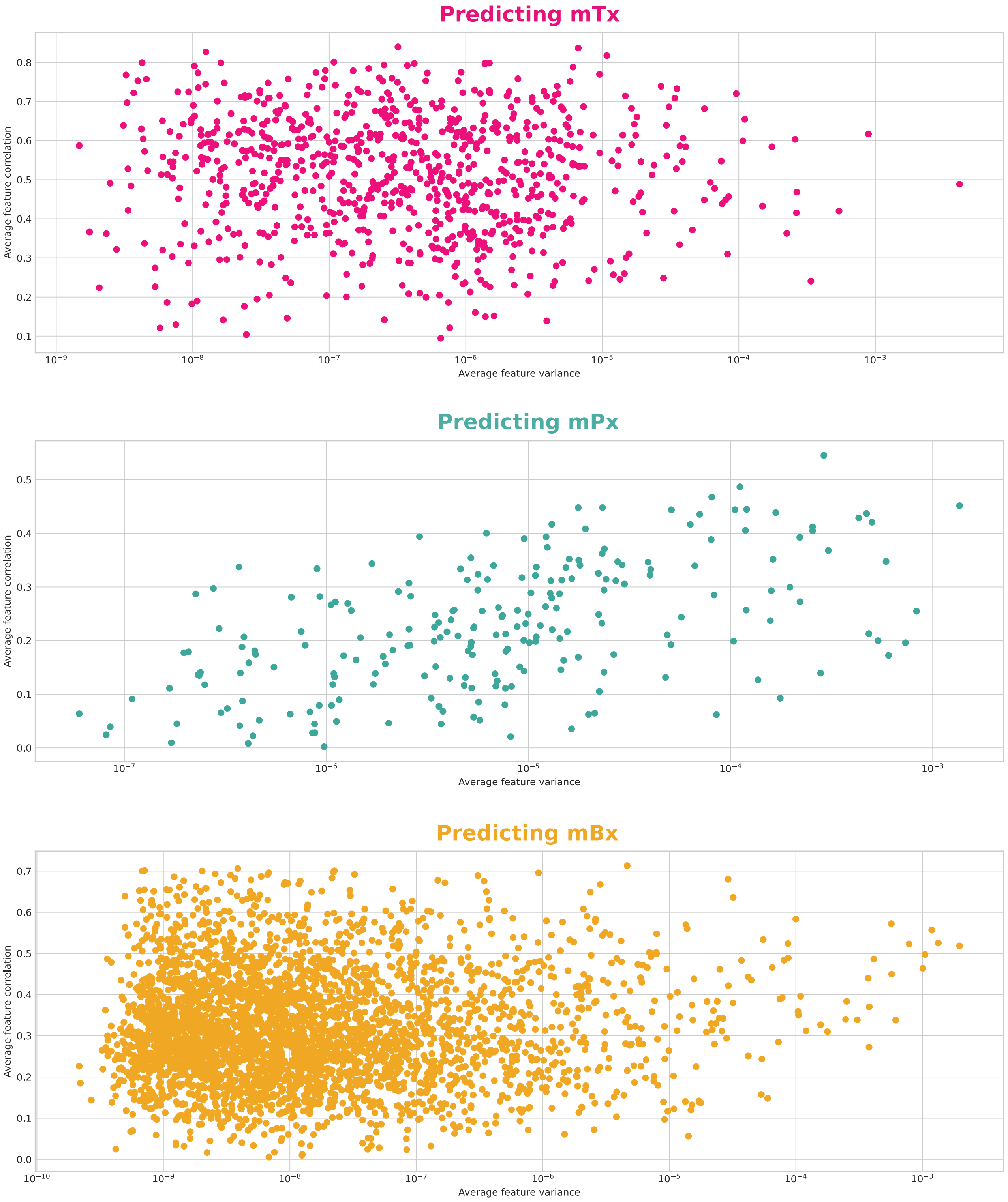


**Figure S3:** Feature variance plotted against feature correlation, for the three output data types. Correlation was computed between predicted features and the ground-truth. Variance and correlation were both computed on test sets, and averaged across dataset partitions and single- and multi-omics input types.

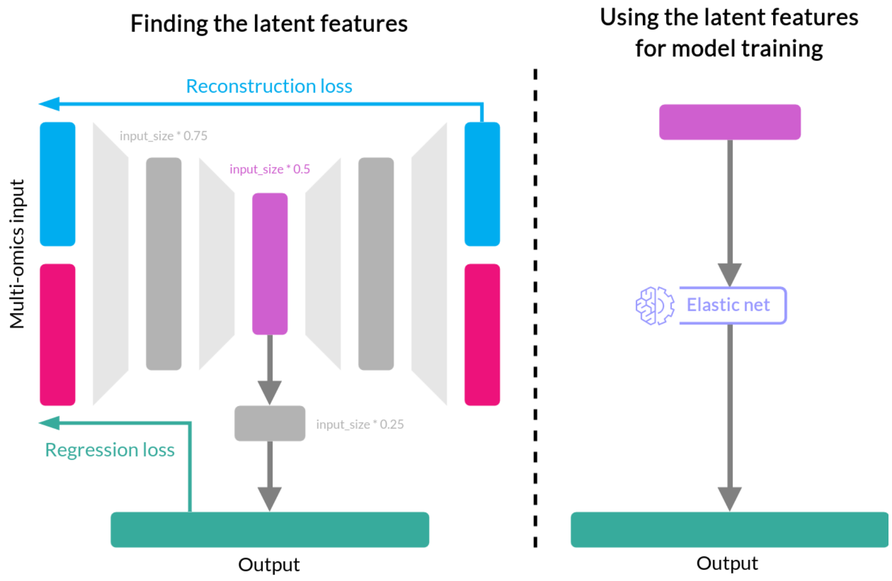


**Figure S4:** Training a multi-omics autoencoder (Supplementary Note A.2) with a combined loss, followed by training an elastic net model (MelonnPan[2]) on the latent features.

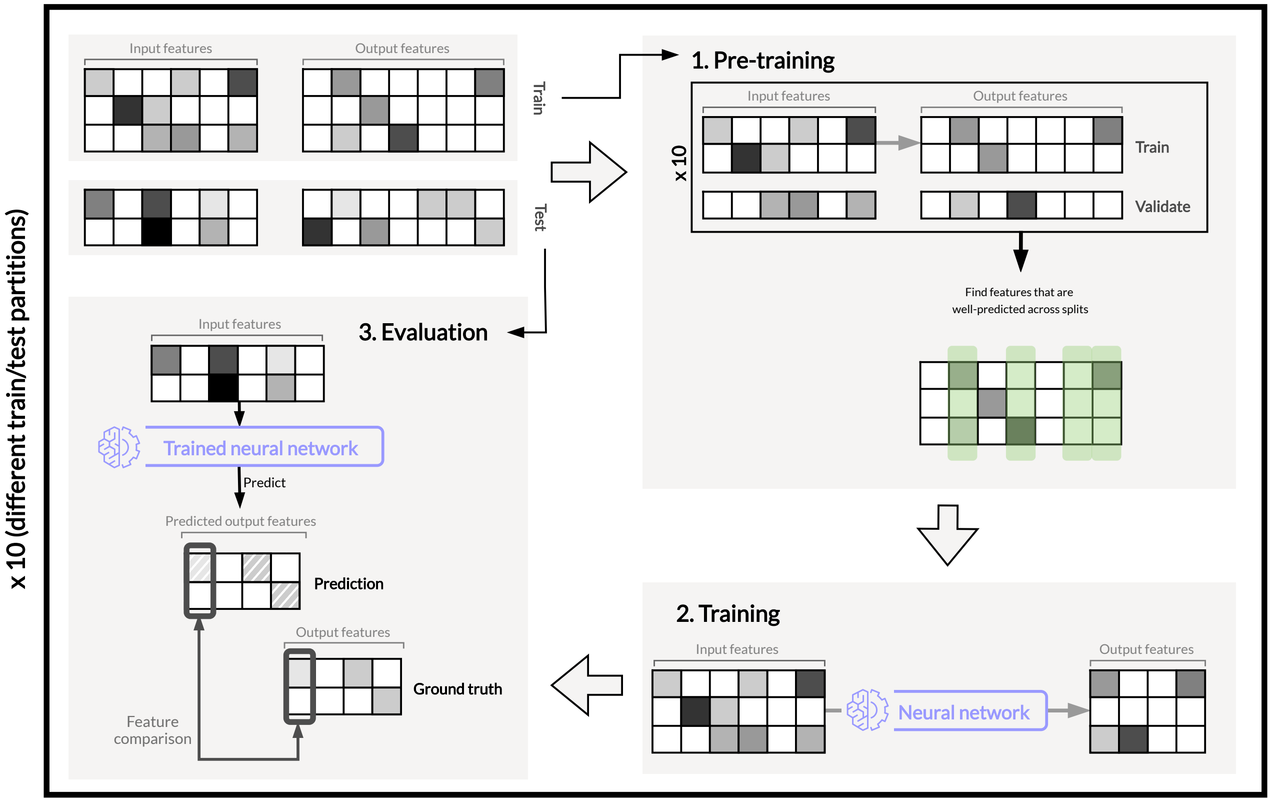


**Figure S5:** Pre-training for feature selection, performed on cross-validation folds. Selected features are subsequently used to train a neural network, as the one described in Supplementary Note A.1.
